# Effects of Lamina-Chromatin Attachment on Super Long-Range Chromatin Interactions

**DOI:** 10.1101/2025.02.13.638183

**Authors:** Pourya Delafrouz, Hammad Farooq, Lin Du, Ao Ma, Jie Liang

## Abstract

The interactions between chromatin and lamin proteins localized on the nuclear envelope play a crucial role in the three-dimensional (3D) organization of the genome. This study investigates the influence of lamin associated domains (LADs) on genome organization at the chromosome level using 3D polymer models of mouse embryonic fibroblasts (MEFs) and embryonic stem cells (mESCs). By integrating genome-wide LAD maps from DamID assays, we simulated chromatin conformations with and without LAD attachment to the nuclear envelope. Our results show that incorporating LAD-lamin interactions yields a radial chromatin distribution consistent with experimental observations. Moreover, LAD-lamin interactions induce significant super long-range chromatin contacts across distant genomic regions. These findings suggest two distinct mechanisms driving induction of chromatin interactions by LAD-lamin attachment.

## 1. Introduction

The nuclei of eukaryotic cells at interphase are well organized. Individual chromosomes occupy non-random territories [1, 2] and maintain a hierarchy of structures at different scales, including compartments [3], topologically associating domains (TADs), and chromatin loops [4].

A major factor in determining the nuclear and genome structures is the interactions between chromatin and nuclear lamina, where lamin proteins are in contact with specific regions of the chromosomes [5]. These regions, called the lamina-associated domains (LADs) [6, 7], have been identified through methods such as DamID measurement, where adenine methylated DNA regions due to contact with Lamin B1 protein in the nuclear lamina are identified [8]. LADs are found to be of size from 0.1 to 10 Mb, can cover about 30–40% of the genome, and have lower gene density [7]. The interactions between LADs and lamin are stochastic, as only a fraction of the LADs are in contact with lamin in a particular cell [9]. LAD-lamin attachment generally provide a repressive environment for gene expression [10, 11] and they are enriched in B compartment [12, 13] Computational studies have shed light on the relationships between LAD-lamin attachment and the reorganization of genome [14, 15, 16, 17, 18, 19, 20, 21, 22, 23, 24, 25, 26, 27]. These Studies have underscored the important roles of lamin-B and B-B compartment interactions in establishing nuclear architecture and the radial distributions of genomic compartments, and by integrating LAD-lamin attachment with other experimental data such as Hi-C, 3D-FISH, and SPRITE, lamin association frequencies can be accurately reproduced [23, 28, 29]. While some studies have focused on the broader aspects of genome organization without detailed considerations of the significance of LADs[28, 30, 14], others have examined more specific phenomena. The roles of LAD-lamin attachment in reorganization of nuclear architecture have been investigated in studies based on phase-field models [24, 26, 27]. Additionally, polymer model studies have elucidated the impact of the strength of LAD-lamin attachment and heterochromatin distribution on the formation of chromosome territories and the length distribution of LADs [22, 23]. Integration of data from Hi-C and LAD information has further advanced methodologies such as Chrom3D, enabling the investigation of radial positioning dynamics of various genomic elements including LADs, topologically associating domains (TADs), and genes [14, 31].

However, current efforts on studying the effects of LAD-lamin attachment are limited to smaller genomes such as *Drosophila* [30], or one or a few chromosomes of mammalian genomes [23, 19]. In addition, thorough sampling of diverse chromosome conformations independent of initial conditions to account for the extraordinary structural heterogeneity of chromosome remains challenging. As a result, important aspects of global long-range effects of LAD-lamin attachment are unknown. We do not know whether chromatin interactions as specific patterns detected in Hi-C studies can arise from distant LAD-lamin attachment.

In this study, we develop a 3D poylmer model and investigate the long-range effects of LAD-lamin attachment on genome organization. We use Mouse Embryonic Fibroblast (MEF) and mouse Embryonic Stem Cell (mESC) as our model systems, since LAD-lamin attachment experience significant changes when embryonic cells develop into differentiated cells [32]. Our study is among the few where 3D conformations of full diploid mammalian genome are considered [28, 15]. Further, our model is well suited to account for the extraordinary heterogeneity in the 3D conformations of the diploid genome to quantitatively describe the geometric properties of chromosomes in single-cells and in populations. As our model is based on analysis of 20,000 single-cell diploid genome conformations, each independently generated using the CHROMATIX deep sampling techniques [33]. This overcomes limitations in sampling at mixing due to finite simulation time in molecular dynamics based studies, where harvested conformations in a simulation trajectories are likely correlated due to the limited time scale of the simulations.

A remarkable finding is that LAD-lamin attachment induces chromatin interactions broadly across all chromosomes, even in regions distant from LAD regions. Furthermore, the induced chromatin interactions are overwhelmingly between inactive chromatin regions, suggesting their roles in suppressing gene expression. In addition, we identified a small number of specific chromatin interactions that reproduce Hi-C interaction patterns of the whole genome, with their numbers inversely related to the size of the induced chromatin interactions.

## 2. Methods

### 2.1 A diploid genomic polymer model incorporating LAD-lamin attachment

Previous polymer models of chromatin for cells have focused on intra- and inter-chromosomal interactions [34, 17, 23, 33, 36]. Several studies have also incorporated lamina-chromatin interactions, either for a selected set of chromosomes [23, 37] or for the whole genome diploid [38, 15, 14].

Ensemble models of diploid whole genome is challenging, because the extraordinary heterogeneity of 3D genome structures requires the generation of a large ensemble of independent genome conformations from a well-defined distribution. For this purpose, we employ a deep sampling approach based on sequential Monte Carlo chain growth to ensure thorough sampling and to avoid local minima often encountered in molecular dynamics simulations [36, 33, 39].

#### 2.1.1 Polymer model of chromosome and LADs

To model the full diploid mouse genome of 2*×*20 chromosomes of 2,723 Mb in MEF and mESC cells, we construct a coarse-grained polymer model, where each polymer bead represents 500 kb and has a diameter of 188 nm based on consideration of the DNA fiber density of 11 nm [40]. A total of 2*×*5, 278 = 10, 556 beads are used to represent the diploid genome. The nuclei radius of MEF is taken as 11.90 *µ*M [41]. The LAD regions on the chromosomes of MEF and mESC cells are identified by DamID [32, 7]. We use a total of 2*×*1,200 and 2*×*1,050 beads to account for the LAD regions for MEF and mESC, respectively.

We develop three different classes of 3D genome models.

- The *NoLAD ensemble* consists of 3D genome conformations, where the 2*×*20 chromosomes are modeled as 40 self-avoiding random walks of appropriate length with excluded volume, all confined within the nuclear envelope (Fig 1B). This class of model captures essential properties of random polymers and can account for scaling rules observed from Hi-C measurements [42]. With a few additional landmarks, they can also reproduce Hi-C heatmaps of the budding yeast [39].
- The *LAD ensemble* consists of 3D genome conformations with LAD beads distinguished from others. Region of LAD beads are obtained from DamID mapping (Fig 1A). Each 3D genome conformation has a total of 2*×*1,200 = 2,400 LAD beads among the 2*×*5,278 polymer beads. As 30% of LAD regions are found to be attached to the lamina [43] and such attachment is stochastic [44, 45], 800 beads out of the 2,400 LAD beads in each conformation are set to be attached to the nuclear envelope in the ensembles for MEF cells (700 out of 2,100 LAD beads are attached to the envelope for mESC cells). Here beads within the distance threshold of 0.6 *µ*m to the nuclear envelope are considered to be attached.
- The *Hi-C ensembles* consists of 3D genome conformations that reproduce Hi-C frequencies. As recent studies showed that a set of sparse specific polymer interactions derived from Hi-C frequencies are sufficient to fold chromatin loci in *Drosophila* [36] as well as in human cells [46], we also construct *Hi-C ensembles* for each chromosome, where a set of specific polymer interactions accounting to ~15% of Hi-C interactions are used to generate ensembles of single-cell chromosome conformations [21].

**Figure 1:**
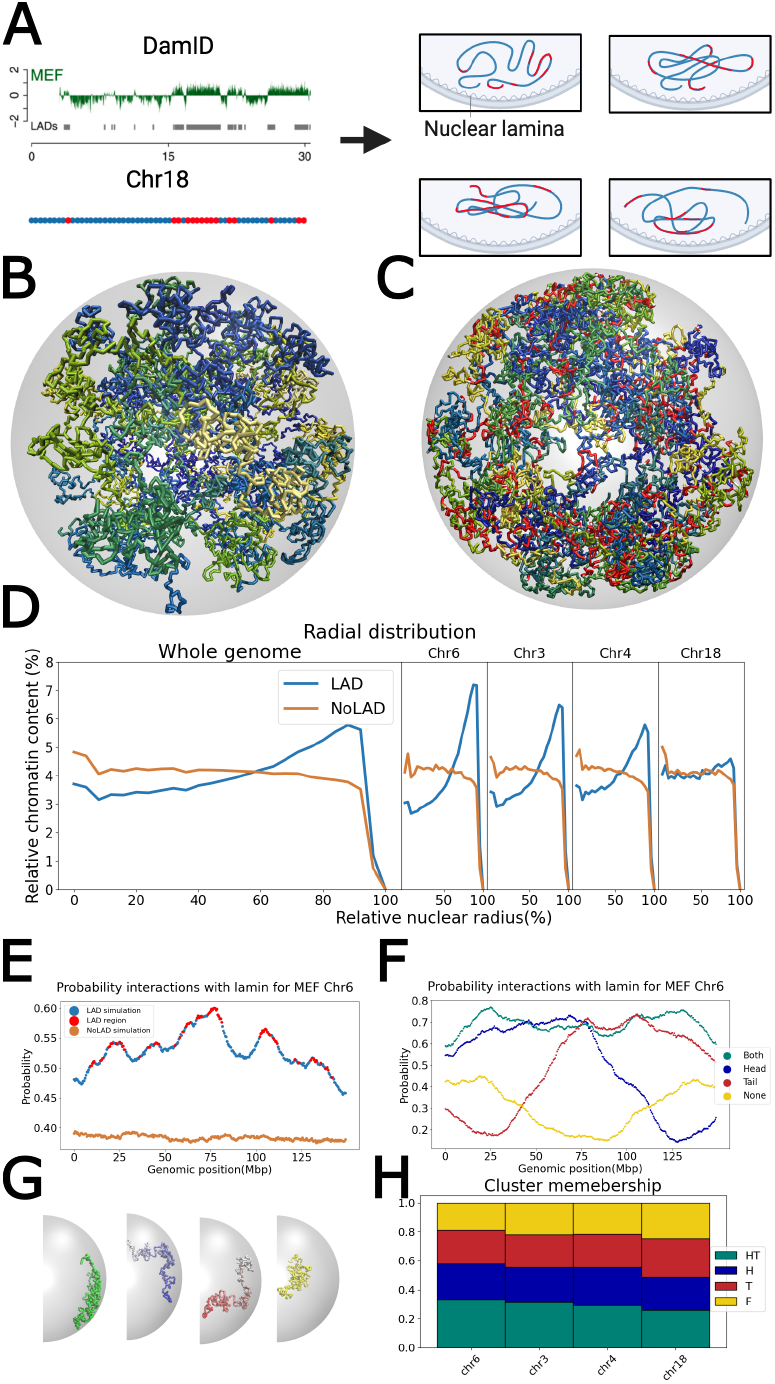
Polymer model of the diploid genome of mouse embryonic fibroblast (MEF) cells. A) Using DamID measurement [32], the LAD regions, depicted in red, and the remaining regions, depicted in blue, are derived. Creating conformations by considering the portions of LAD regions close to the nuclear lamina in each conformation. B) An example of the 3D conformation of the diploid genome of the *NoLAD* ensemble, with each chromosome colored differently. C) An example of 3D conformations *LAD* ensemble, with LAD regions are shown in red. D) The radial distribution of chromatin density of the diploid genome. With LAD-lamina attachment, chromatin density in the *LAD* ensemble increases along the radial direction towards the peripheral, while, the random polymer ensemble exhibits a generally flat distribution. The radial distributions of chromatin in Chr 6, 3, 4, and 18 in decreasing order of proportion of LAD (31%, 26%, 21%, and 11%) are shown. The peak density near the peripheral is reduced as the fraction of the LAD region decreases. E) The probability of regions along Chr 6 to be attached to the nuclear envelope. For the LAD ensemble, LAD regions have the highest probabilities of attachment. In contrast, the random ensemble has a uniformly lower probability for nuclear envelope attachment. F) The average probability of contact with nuclear envelope of the four distinct conformation classes for Chr 6. G) Illustration of the four distinct conformation classes. Those proximate to lamin in both head and tail regions are in green, those near lamin in the head region only are in blue, those near lamin in the tail region only are in red, and those not close to either the tail or head region are in yellow. H) Cluster size of different conformation types for chromosomes in decreasing order of LAD proportions. As the LAD size decreases, the fraction of conformations closer to the lamin also decreases.

Each of the conformations is generated independently through chain growth employing deep sampling techniques [33], allowing thorough sampling of highly heterogeneous 3D genome structure. Altogether, we have an ensemble of a total of 20,000 3D genome conformations for each of the *LAD* and the *NoLAD* ensemble.

## 3. Results

### 3.1 Simulation reproduces radial distributions of chromatin density and characterizes heterogeneity of LAD-lamina contact frequencies

Taking the ensembles of 20,000 single cell conformations of *NoLAD* (Fig 1B) and *LAD* (Fig 1C), we examine how the attachment of LAD to lamin affects various cellular properties across different scales for MEF cells. We first calculate chromatin density at varying distances from the nucleus center. With LADs, the chromatin density of the diploid genome (Fig 1D, left panel) increases from the nucleus center towards the cell periphery, peaking at 90% of nuclear radius. Conversely, in the absence of LAD-lamin attachment, chromatin density peaks at the nuclear center and decreases slightly towards the periphery, resulting in an overall flat distribution from 0 to 95% of nuclear radius. These results resemble the radial chromatin distributions observed in *Drosophila*, both through fluorescence measurements [47] and simulations [30]. Moreover, the impact of lamina attachment is directly proportional to the fraction of LAD regions on each chromosome (Fig 1D, four right panels). The most pronounced effects are observed on Chr 6, which has the largest fraction of LAD regions (31%), while minimal effects are seen on Chr 18, which has the smallest fraction of LAD regions (11%). Details on other chromosomes can be found in Supplementary Fig S1.

Additionally, we examine the likelihood of different regions of chromosome to be in close proximity to the nuclear envelope. As illustrated for Chr 6 in MEF (Fig 1E), most LAD regions are at local probability peaks to be near the nuclear envelope, as anticipated. The height of peak is correlated with the lengths of the LAD regions. The probability of envelope attachment is the highest of 0.6 at genomic location of 77 Mbp. At this location, Chr 6 also has its most extensive LAD region, spanning 7.5 Mbp.

Chromosomes in individual cells can adopt divers configurations. To quantify how LAD-lamin attachment differ among the conformations, we classify the chromosome configurations broadly into four distinct clusters of conformations based on their patterns of LAD-lamin attachment, as seen in Fig 1F for Chr 6 and illustrated in Fig 1G:

- “HT” head-and-tail configurations (green), where the entire chromosome region is in close proximity to the nuclear periphery.
- “H” head configurations (blue), where the head portion of the chromosome is in close proximity the periphery while the tail extends further away.
- “T” tail configurations (red), where the tail of the chromosome is in close proximity with the nuclear periphery while head extends further away.
- “F” free configurations (yellow), where the chromosome is completely distant from the nuclear periphery.

Such heterogeneity in LAD-lamin attachment is seen in all chromosomes, as all four classes of cluster are present for all chromosomes (Fig 1H). There are, however, patterns in cluster populations across different chromosomes. As shown in Fig 1H, Chr 6 which has the largest LAD region, its “HT” cluster has a larger population compared to Chr 18, which has the smallest LAD region. In contrast, Chr 18 has the largest population for “F” cluster compared to all other chromosomes (see Supplementary Fig S2 for all chromosomes in MEF and mESC cells). Our results show that the largest LAD portion for chromosome forced more conformations closer to the nuclear envelope (“HT”); however, we still have conformations distant from the nuclear periphery (“F”), but they are less common.

These findings demonstrate that our model is consistent with the results from DamID assay of MEF and mESC and replicates the radial distribution observed in experiments [48]. Moreover, it quantitatively reveals the heterogeneity in LAD-lamin attachment [43, 9].

### 3.2 LAD-lamin attachment induce Hi-C chromatin interactions broadly

#### 3.2.1 LAD-lamin attachment induces significant amount of Hi-C chromatin interactions

While the overall effects of LAD-lamin attachment on radial distribution of chromatin are well known [7], whether Hi-C patterns of chromatin interactions can arise from LAD-lamin attachment is unknown.

We use the *LAD* and *NoLAD* ensembles to investigate this issue. We first remove the random polymer effects by subtracting simulated Hi-C interactions of the *NoLAD* ensemble from both experimental Hi-C and simulated Hi-C of the *LAD* ensemble (Fig 2A). We then identify regions in the experimental Hi-C map that are strongly correlated with those of the LAD model after adjustment of multiple hypothesis testing [49].

**Figure 2:**
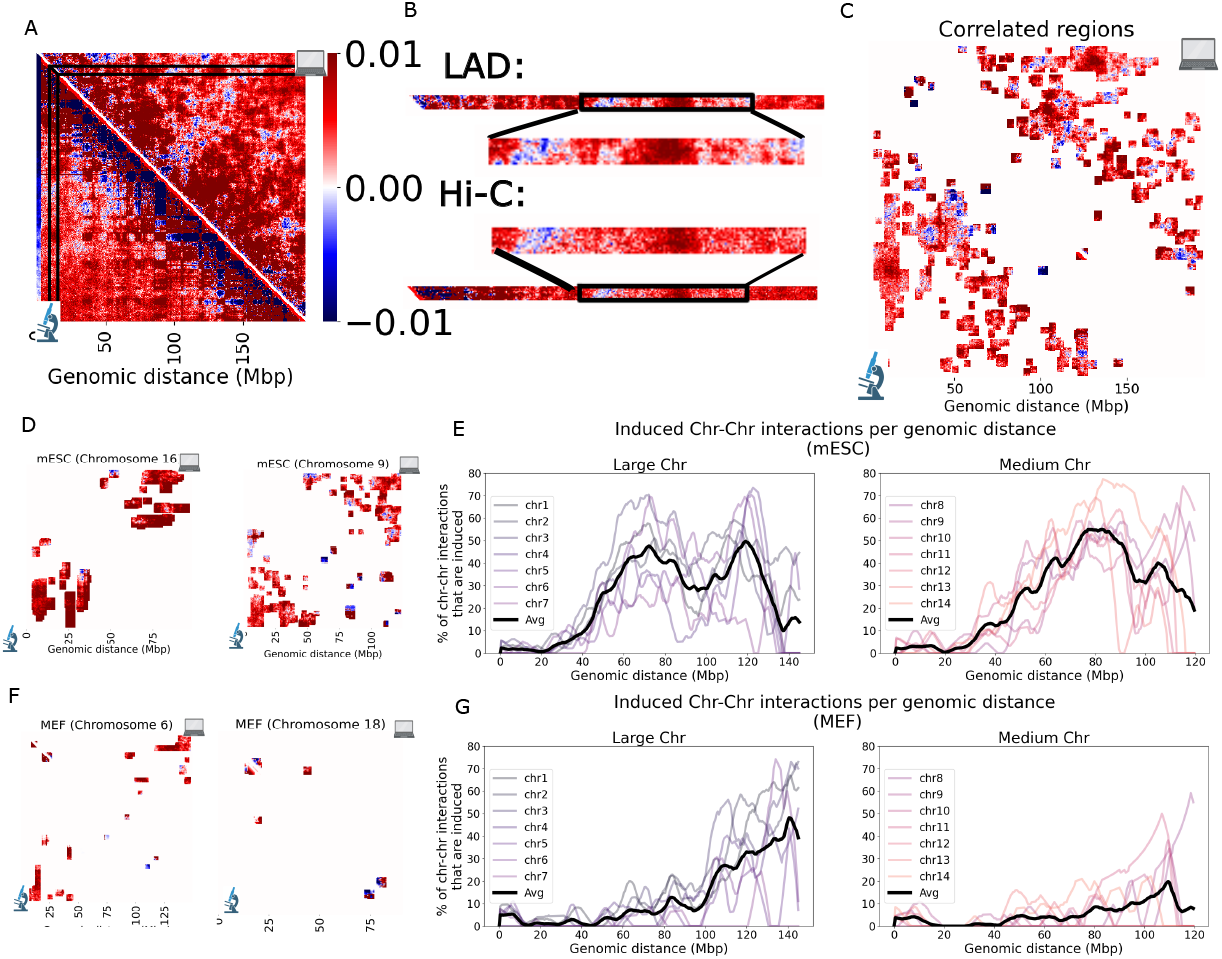
LAD-induced Hi-C chromatin interactions. A) Identification of correlated regions in Hi-C maps. The contact frequencies from both the Hi-C map and the *LAD* model are shown, with the *NoLAD* model subtracted to remove the random polymer effect and identify the correlated regions. Right panel shows a region of the chromosome interacting with the rest of the chromosome in both the *LAD* model and the Hi-C map. It identifies correlated regions, interactions between Chr1:88.5-164 Mbp and Chr1:9.5-15.5 Mbp as correlated in both the Hi-C map and the *LAD* model. B) Correlated region between the *LAD* model and experimental Hi-C map for mESC Chr1. 23% of Hi-C area is correlated. C) LAD-induced Hi-C chromatin interactions in mESC for Chrs 16 and 9 with most and least LAD sizes. The majority of correlated regions are observed to be further from the main diagonal showing long-range effects of LAD on chromatin organization. D) Distribution of fraction of chromatin interactions which are induced by LAD-lamin interactions across various genomic distances for mESC across large and medium chromosomes (100% shows all chromatin interactions with a specific genomic sepration distance). In large chromosomes, half of Hi-C chromatin interactions at genomic separation of 72.5 Mbp and 120.0 Mbp are induced by LAD-lamin attachment. The pattern for medium chromosomes is similar with larger chromosomes. It shows the long-range effects of LAD-lamin attachment for both large and medium chromosomes in mESC. E) LAD-induced Hi-C chromatin interactions in MEF for Chrs 6 and 18 with most and least LAD sizes. Owing to the condensed structure of MEF, the majority of correlated regions are observed to be positioned between the two ends of the chromosomes. F) Distribution of fraction of chromatin interactions which are induced by LAD-lamin interactions across various genomic distances for MEF across large and medium chromosomes (100% shows all chromatin interactions with a specific genomic sepration distance). In large chromosomes in MEF, half of Hi-C chromatin interactions at genomic separation of 140.5 Mbp are induced by LAD-lamin attachment. However, for medium chromsomes in MEF, the amount of LAD-induced Hi-C chromatin interactions is very small (Supplementary Fig S5 for small chromosomes). It shows that effects of LAD-lamin attachment is more long-range for MEF compared to mESC. The difference between MEF and mESC in the distribution of chromatin interactions induced by LAD-lamin interactions is related to the more closed chromatin structure of MEF.

As an example, Fig 2B shows Hi-C interactions between the region of Chr1:9.5-15.5 Mbp and the rest of Chr 1 in mESC cells in both the LAD model and the experimentally measured Hi-C study after removing random polymer effects. The interaction region between Chr1:9.5-15.5Mbp and Chr1:88.5-164.0 Mbp exhibit a strong correlation of 0.74 between *LAD* simulated and measured Hi-C maps.

There are numerous such strongly correlated regions in the Hi-C maps. Chromatin-chromatin interactions as measured in the Hi-C study are likely induced by LAD-lamin interactions. Such induced regions appear across the whole chromosome, covering 23% of all Hi-C regions in Chr 1 (Fig 2C). that are strongly correlated with that of the *LAD* model.

The LAD-induced Hi-C chromatin interactions are a general phenomenon, as they are found across different chromosomes. Chromosomes with large or small LAD regions can all have significant amount of LAD-induced Hi-C chromatin interactions. As an example, Chr 16 and Chr 9, with 40% and 8% covered by LAD, respectively, both exhibit significant amount of LAD-induced Hi-C chromatin interactions (16% and 17%, respectively. Fig 2D). Other chromosome with a small fraction of LAD-lamin attachment are also found to exhibit a significant amount of induced Hi-C chromatin interactions. Chr 1/2/6 with 25%/23%/24% covered by LAD have 23%/26%/25% of the chromatin-chromatin interactions induced by LAD-lamin interactions (Supplementary Fig S3).

#### 3.2.2 LAD-lamin attachment induced chromatin interactions are super-long range

The effects of LAD-lamin attachment on Hi-C pattern are super long-range in nature. Across the genome, 41% of the 198,910 chromatin interactions in mESC cells between regions more than 60 Mbp apart (at 500 kb resolution) are driven by LAD-lamin attachment. For all large chromosomes (Chr 1–7 with length 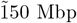), LAD-lamin attachment significantly alter chromatin-chromatin contacts. When the interacting regions are 120.0 Mbp apart, about half of all Hi-C contacts are LAD-induced (Fig 2E). At 72.0 Mbp separation, again about half of all Hi-C contacts are induced by LAD-lamin attachment. In contrast, less than 5% of Hi-C contacts between regions that are *<*20 Mbp apart are induced by LAD-lamin attachment. For chromosome of medium size (Chr 8–14, from 120 Mbp to 150 Mbp), similar behavior are observed with only minor differences. Specifically, when the chromatin regions are 80 Mbp apart, 54% of all Hi-C measured chromatin-chromatin contacts are induced by LAD-lamin attachment. These results show that for both large and medium sized chromosomes, LAD-lamin attachment is a strong determinant of super long range (≥ 72 Mbp) Hi-C chromatin contacts.

#### 3.2.3 LAD-lamin induced interactions are less prominent at later developmental stages

LAD induced Hi-C chromatin interactions can occur in cells at different stages of development, as effects of LAD-lamin attachment are also seen in mouse embryonic fibroblast (MEF), albeit to a less extent. In MEF, Chr 6 has the largest fraction of LADs (31%) and about 5% of the Hi-C regions are LAD induced. Chr 18 has the smallest fraction of LADs (12%) and only 1% of the Hi-C regions are LAD induced (Fig 2F, Supplementary Fig S4). Among large chromosomes (Fig 2G), strong effects on Hi-C chromatin interactions are seen when chromatin regions with a separation ≥100 Mbp, peaking at 140.5 Mbp. Among all medium chromosomes, LAD induced Hi-C contacts are negligible. LAD-lamin attachment affects fewer chromatin interactions in MEF compared to mESC. Additionally, it affects regions that are farthest from each other (Fig 2E).

The differences in LAD induced effects are likely due to the more developed nucleus of MEF cells. During differentiation from mESC to MEF, there is significant amount of increase of heterochromatin [50]. This is due to numerous chromatin remodeling events occurring during this period as machinery of gene regulation matures. As heterochromatin regions are tightly packed and are associated with repression of transcription, increase in heterochromatin restricts the open regions and reduces the mobility of chromatin in the nucleus. As a result, the ability of LAD-lamin attachment to induce Hi-C chromatin interactions is severely hampered. In summary, our physical models uncovered that significant amount of chromatin contacts captured in Hi-C studies are induced by LAD-lamin attachment. In mESC, even chromosomes with a small fraction of LAD regions can exhibit a large amount of LAD induced Hi-C chromatin interactions. The LAD-lamin attachment effectively promotes super long-range interactions. The impact of LAD-lamin attachment in inducing chromatin-chromatin interaction is reduced in mESC, due to the more structured cell nucleus and increased heterochromatin.

#### 3.2.4 Mechanism of induced Hi-C chromatin interactions by LAD-lamin attachment for induced Hi-C interactions between LAD and Non-LAD regions

It is important to examine the mechanism of induction effects by LAD-lamin attachment. Further examination of LAD-induced chromatin contacts, about half of them occur outside the LAD regions. That is ~ 12% Hi-C chromatin interactions in Chr 1 are between Non-LAD regions dictated by LAD domains (Fig 3A). LAD-induced Hi-C chromatin interactions can occur through two mechanisms. *LAD Crowding* happens when limited free space in the peripheral of the nucleus pushes chromatin near the nuclear lamina to be proximal with LAD regions leading to Hi-C contacts(Fig 3B, top panel). *LAD Anchoring* occurs when two LAD regions interacting with the nuclear lamina pull adjacent chromatin closer together (Fig 3B, bottom panel).

**Figure 3:**
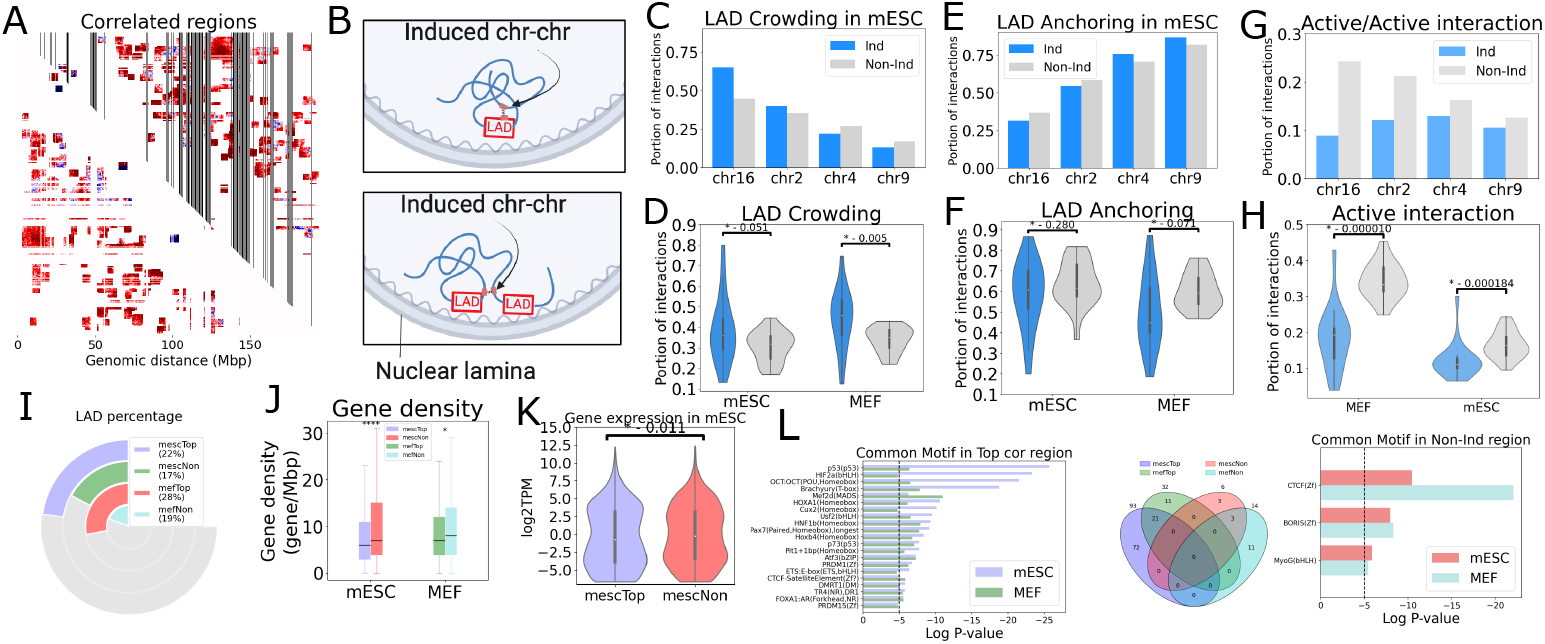
LAD-induced Hi-C chromatin interactions characteristic. A) Correlated regions for mESC Chr 1 with LAD regions showing as black strip. The correlated regions are beyond LAD regions with 48% of correlated regions outside of LAD regions. B) Two mechanism of LAD-lamin attachment induce Hi-C chromatin interactions (pink): Top panel) Due to nuclear volume confinement, chromatin interacts with LAD regions that are attached to the nuclear lamina (*LAD Crowding*). Bottom panel) Neighboring regions of two LADs interact with the nuclear lamina near each other (*LAD Anchoring*). C) *LAD Crowding* can quantify by LAD/Non-LAD interactions within induced Hi-C chromatin interactions of mESC chromosomes with different amount of LAD regions, for Chr 16 (39% LAD, highest) to Chr 9 (8% LAD, lowest). As LAD region decreases, the difference in *LAD Crowding* diminishes, from 20% (Chr 16) to −4% (Chr 9). D) *LAD Crowding* in mESC and MEF across all 20 chromosomes. *LAD Crowding* differ significantly in both MEF and mESC between induced and Non-induced regions. It shows the significance of *LAD Crowding* mechanism as one of mechanism for inducing Hi-C chromatin interactions. E) *LAD anchoring*, involving Non-LAD/Non-LAD interactions, shows no significant difference across mESC chromosomes with varying amounts of LAD regions, changing from −5% (Chr 16) to 5% (Chr 9). F) *LAD anchoring* are similar between induced and Non-induced regions for both MEF and mESC. It shows the magnitude of *LAD anchoring* in induced regions is not significantly different from that in non-induced regions. G) Activity of regions in *LAD anchoring* for mESC with different chromosomes and LAD sizes. For chromosomes with the most LAD, inactive regions are predominantly in induced Hi-C chromatin interactions. H) Fraction of Active interactions in *LAD anchoring* across all 20 chromosomes. Non-induced Hi-C chromatin interactions contain significantly more Active interactions compared to induced interactions for both mESC and MEF, suggesting in *LAD Anchoring* mechanism force the inactive regions interact with each other. I) Percentage of LAD for “Top” and “Non”-induced regions. Although induced regions have higher LAD percentages, but it is still small portion of them are LAD regions (22% for mESC and 28% for MEF). J) Gene density through for “Top” and “Non”-induced regions. “Non”-induced regions exhibit higher gene density. K) Gene expression “Top” and “Non”-induced regions in mESC. The data reveals elevated log2TPM values for “Non”-induced regions, indicating higher gene expression. L) Common motifs between “Top” and “Non”-induced regions in mESC and MEF. Notably, there is no shared motif between mescNon vs mefTop and mefNon vs mescTop. The result indicates high level of CTCF and Boris motifs in “Non”-induced regions, associated with chromatin structure for both MEF and mESC. Additionally, “Top” regions exhibit higher levels of p53 and Mef2d motifs, with previous studies suggesting their connection to LAD.

*LAD Crowding* can be quantified by calculating the ratio of LAD/Non-LAD interactions to the total interactions (LAD/LAD, Non-LAD/Non-LAD, and LAD/Non-LAD). By comparing this ratio between the induced and non-induced regions (used as a control), we can observe how this mechanism operates. Chr 16, which has the highest number of LAD regions in mESC, shows an increased LAD/Non-LAD interaction ratio in induced Hi-C chromatin interactions compared to non-induced interactions (Fig 3C). Chr 2 also displays an elevated ratio. In contrast, Chr 9, which contains the fewest LAD regions, exhibits a similar LAD/Non-LAD interaction ratio in both induced and non-induced interactions. Chr 4 also shows similiar behavior as Chr 9. These results suggest that the degree of difference in LAD/Non-LAD interactions between induced and non-induced regions correlates with LAD size, highlighting *LAD Crowding* as a potential mechanism.

Overall, LAD-induced Hi-C chromatin interactions across all chromosomes show an increase in the LAD/Non-LAD interaction ratio. The average LAD/Non-LAD interaction ratio in induced regions is 38% in mESC and 44% in MEF, compared to 31% and 34% in non-induced regions, respectively (Fig 3D). These values are significantly lower in non-induced regions. Higher ratio of LAD/Non-LAD interactions in induced region suggest that *LAD Crowding* is one of the mechanisms driving LAD-induced Hi-C chromatin interactions, particularly in chromosomes with larger LAD regions.

The other mechanism, *LAD Anchoring*, can be quantified by calculating the ratio of Non-LAD/Non-LAD interactions to the total interactions. For these interactions, the difference between induced and non-induced regions is minimal, regardless of the chromosome’s LAD size as shown for Chrs 16, 2, 4, and 9 (Fig 3E). Furthermore, across all chromosomes in both MEF and mESC, there is little variation in Non-LAD/Non-LAD interactions between induced and non-induced regions (Fig 3F). Our results show that the *LAD Anchoring* mechanism is similar in the amount for induced and non-induced regions.

The underlying genomic activities of *LAD Anchoring* mechanism differs significantly between induced and non-induced regions. Here, we analyze Hi-C data to identify A/B compartments and use H3K27ac histone modification markers to pinpoint active genomic regions in Non-LAD/Non-LAD interactions. As seen Fig 3G, Chrs 16, 2, 4, and 9 in mESC cells all have less Active/Active genome interactions in induced regions than non-induced regions. This difference between induced and non-induced is the largest in Chr 16, which has the most LAD regions, compared to Chr 9, which has the least. Our results across chromosomes, the fraction of inactive interactions in induced region is correlated with LAD size.

Conversely, the fraction of active genomic interactions for Non-LAD/Non-LAD regions is higher in noninduced regions compared to induced regions (Fig 3H). The average ratio of active genome interactions in Non-LAD/Non-LAD interactions for non-induced regions and induced regions is 32.6% (16.4%) and 18.8% (12.1%) for MEF (mESC), respectively. The higher ratio of inactive interactions in Non-LAD/Non-LAD regions, representing *LAD Anchoring*, in induced regions, suggests that the *LAD Anchoring* mechanism predominantly affects inactive regions.

Our findings suggest that induced regions arise from two distinct scenarios: *LAD Crowding*, LAD regions interact with the nuclear lamina, and due to nuclear volume constraints, Non-LAD regions are forced to interact with LAD regions. *LAD Anchoring*, two LAD regions may force neighboring Non-LAD regions to interact and those regions are inactive.

#### 3.2.5 Regions exhibiting enhanced induction of interactions

We examine regions exhibiting the highest degrees of induced interactions. Based on the number of interactions in induced regions for each 500 kb genomic region, we categorize the top five regions as “Top” and the bottom five regions as “Non”. A significant proportion of non-LAD regions are found in the “Top”-induced regions (78% in mESC and 72% in MEF, Fig 3I), indicating that LAD-lamin interactions involve extensively in Non-LAD regions in the strongly induce regions. Our results show that induced chromatin interactions extend beyond LAD.

Further, the “Top”-induced regions have lower gene density than “Non”-induced regions (Fig 3J). The average gene density in “Top”-induced regions for mESC (MEF) cells is 7.96 (9.61) genes/Mbp, compared to 10.28 (10.56) genes/Mbp in “Non”-induced regions. In addition, genes in “Top”-induced regions exhibit lower expression levels compared to “Non”-induced regions, with an average log2TPM of −0.03 for Non-induced regions and −0.3 for Top-induced regions (Fig 3K). Overall, these results show that induced region have low gene density and low gene activities.

The significance of induced regions is further underscored through analysis of motif enrichment. Motifs enriched in both “Non” and “Top” region are identified using HOMER tool [51]. We identify motifs that are not shared between “Top” and “Non” regions in each cell type. The Venn diagram in Fig 3L shows the overlap of motifs among four regions; mescTop, mescNon, mefTop, and mefNon. As we can see, there are no common motifs between pairs of 1) mescTop and mefNon, and 2) mefTop and mescNon. Our results show that “Top” and “Non” regions from different cell types do not share same motif.

However, there are common motifs identified in “Top” and “Non” regions in MEF and mESC. Motifs associated with LAD, such as p53 [52] and Mef2d [53], are enriched in “TOP” regions for both MEF and mESC. While, motifs related to chromosome structures, CTCF and Boris motifs [54], are enriched in “Non” regions. The results suggest that effect of LAD-lamin attachment on genome organization play important roles in repressed genes, p53 and Mef2d and less involve in genome structure motifs.

In summary, these analyses collectively underscore the influence of LAD-lamin attachment on the chromatin interactions within regions characterized by genomic inactivity and absence of defined motifs related to chromatin structures.

#### 3.2.6 Effects of specific interactions on the LAD-induced Hi-C chromatin interactions

As we mentioned, LAD-lamin attachment do not induce Hi-C chromatin interactions at regions enriched with motifs related to chromatin structures. We further investigate other mechanism for genome organization. Further examine of contact frequency map of *LAD* ensemble and experimental Hi-C data shows the absence of topologically associating domain (TAD) structures in *LAD* ensemble (Fig 4A). Furthermore, the induced regions are mostly inactive regions. Therefore, active chromatin interactions play an essential role in the size and position of the induced regions. In order to investigate the effect of chromatin interactions, we find the set of non-random specific interactions (black dots in Fig 4B) for each chromosome [21]. These specific interactions are enough to get single-cell conformations matching with Hi-C. Contact frequency maps of an ensemble of 20,000 conformations made using these specific interactions, *Hi-C* ensemble, for Chr 1 in mESC (Fig 4B) exhibits a Pearson’s correlation coefficient of 0.91 with experimental Hi-C. Supplementary Fig S10 for MEF and Supplementary Fig S11 for mESC contain the contact frequency maps of all chromosomes in the *Hi-C* ensemble. One example of 3D conformations of *Hi-C* ensembles for mESC is shown in Fig 4C.

**Figure 4:**
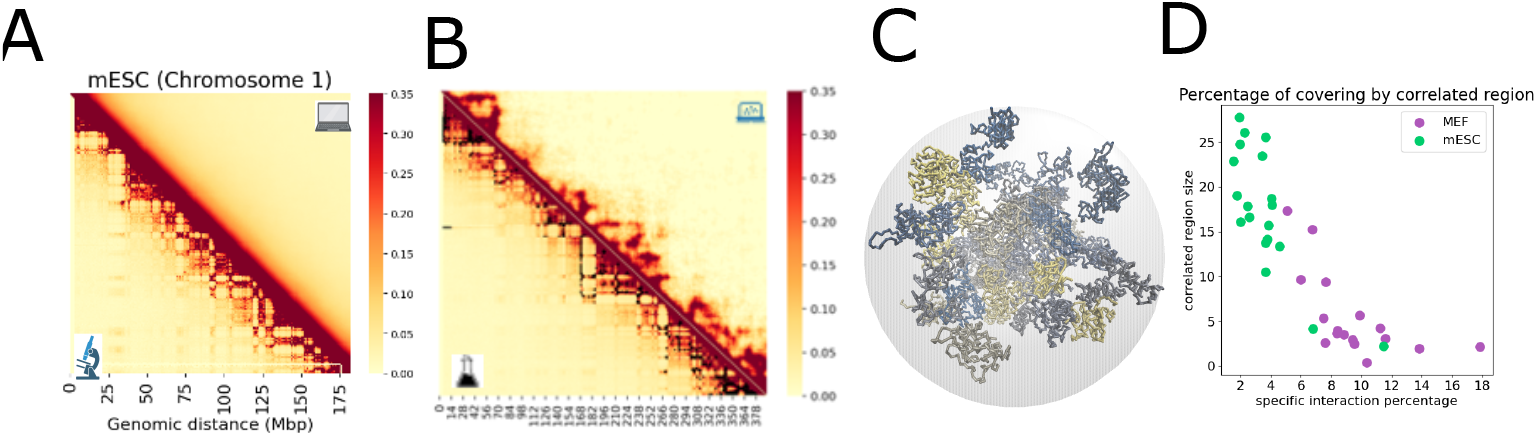
Non-random specific chromatin interactions A) Contact frequency map of LAD simulation and experimental HiC for mESC Chr 1. B) Contact frequency map of simulation using non-random specific interactions (*Hi-C ensemble*, Methods) and experimental HiC for mESC Chr 1 and the Pearson’s correlation is 0.91. Black dots represent non-random specific interactions. C) 3D folded representation of the diploid for mESC. D) Correlation between the percentage of non-random specific interactions and the size of LAD-induced Hi-C chromatin interactions for all chromosomes in mESC and MEF. Suggesting a reverse correlation between the size of LAD-induced Hi-C chromatin interactions and the number of non-random specific interactions for each chromosome.

The percentage of specific interactions is different between chromosomes and cell lines. For MEF, the minimum percentage of specific interactions is Chr 1 with 5.0% of all interactions and the most is for Chr 19 with 17.8%. However, this range is from 1.5% to 11.4% for mESC. Supplementary Fig S8 for MEF and Supplementary Fig S9 for mESC contain the specific interactions for all 20 chromosomes. We see greater number of specific interactions in MEF compared to mESC. Our findings indicate a reverse relationship between the number of specific interactions and the size of LAD-induced Hi-C chromatin interactions (Fig 4D).

We identified that the specific interactions sufficient to form an ensemble of 3D single-cell conformations exhibit a reverse relationship with the size of LAD-induced Hi-C chromatin interactions. The higher number of specific interactions observed in MEF, attributed to its closed chromatin structure, results in an increased number of interactions and smaller LAD-induced Hi-C chromatin interactions.

## 4 Discussion

Our study delves into the intricate landscape of nuclear organization through polymer models, with a particular focus on chromatin lamin interactions. Leveraging a deep sampling approach, we construct a comprehensive ensemble of genome conformations, accounting for the inherent heterogeneity of 3D genome structures. This approach is crucial for capturing the stochastic nature of chromosomal organization and avoiding the entrapment in local energy minima, which is often encountered in traditional molecular dynamics simulations.

A key aspect of the study involves the development of a coarse-grained polymer model that faithfully represents the diploid mouse genome. Each bead in the model corresponds to 500 kb of DNA, allowing for a detailed examination of effects of LAD-lamin attachment. Regions known as lamina-associated domains (LADs), identified through DamID mapping, are incorporated. These LAD regions are known to interact with the nuclear lamina and play a significant role in chromosomal organization and gene regulation.

Using three distinct classes of 3D genome models: the *NoLAD* ensemble, the *LAD* ensemble, and the *Hi-C* ensembles to capture different aspects of chromosomal organization, including the stochastic nature of chromosomal interactions, the spatial distribution of LAD regions, and specific polymer interactions inferred from Hi-C data, respectively.

Chromosomes are categorized into distinct conformational groups based on LAD attachment patterns, with the distribution of interaction frequencies between chromatin and the lamina varying across these clusters. Chromosome with more LADs exhibits more members in the cluster with more frequent interactions with lamin compared to other chromosomes, highlighting the variability in cluster membership across different chromosomes based on LAD size.

First finding of this study is the effect of LAD-lamin attachment on the sub-chromosomal genome organization. A key finding of this study is uncovering regions within Hi-C maps that correlate with contact frequency maps derived from *LAD* simulations. The study employs statistical methods, including quantile normalization and False Discovery Rate (FDR) correction, to identify regions with high correlations. Additionally, the study investigates the biological basis of LAD-induced Hi-C chromatin interactions, revealing dominance of two distinctive mechanisms, *LAD Crowding* and *LAD Anchoring*.

Analyzing strongly induced regions shows that they have lower gene density and lower gene expression levels. Motif analysis unveils regions with less induced chromatin interactions are enriched in chromatin structure motifs such as CTCF and Boris.

Furthermore, we find a small number of specific interactions that they are sufficient for generating singlecell conformations that match experimental Hi-C data. MEF cells exhibit a higher proportion of specific interactions, primarily due to their closed structure, compared to mESC. The reverse relation of number of specific interactions and the size of LAD-induced chromatin interactions are also shown.

Overall, the findings of this study provide a comprehensive understanding of the regulatory mechanisms governing chromosomal organization by LAD-lamin attachment.

## Acknowledgement

This work is supported by NIH grants R03OD036492 and R35GM127084. An award for computer time was provided by the U.S. Department of Energy’s (DOE) Innovative and Novel Computational Impact on Theory and Experiment (INCITE) Program. This research used resources from the Argonne Leadership Computing Facility, a U.S. DOE Office of Sciensce user facility at Argonne National Laboratory, which is supported by the Office of Science of the U.S. DOE under Contract No. DE-AC02-06CH11357.

## Notes

### Competing Interest Statement

The authors have declared no competing interest.

## References

[1] T. Cremer, M. Cremer, Chromosome territories, Cold Spring Harbor perspectives in biology 2 (2010) a003889.

[2] M. Cremer, J. Von Hase, T. Volm, A. Brero, G. Kreth, J. Walter, C. Fischer, I. Solovei, C. Cremer, T. Cremer, Non-random radial higher-order chromatin arrangements in nuclei of diploid human cells, Chromosome research 9 (2001) 541–567.

[3] S. Wang, J.-H. Su, B. J. Beliveau, B. Bintu, J. R. Moffitt, C.-t. Wu, X. Zhuang, Spatial organization of chromatin domains and compartments in single chromosomes, Science 353 (2016) 598–602.

[4] S. Rao, M. Huntley, N. Durand, E. Stamenova, I. Bochkov, J. Robinson, A. Sanborn, I. Machol, A. Omer, E. Lander, E. Aiden, A 3D Map of the Human Genome at Kilobase Resolution Reveals Principles of Chromatin Looping, Cell 159 (2014) 1665–1680. Number: 7.

[5] T. A. Dittmer, T. Misteli, The lamin protein family, Genome biology 12 (2011) 1–14.

[6] A. Buchwalter, J. M. Kaneshiro, M. W. Hetzer, Coaching from the sidelines: the nuclear periphery in genome regulation, Nature Reviews Genetics 20 (2019) 39–50.

[7] L. Guelen, L. Pagie, E. Brasset, W. Meuleman, M. B. Faza, W. Talhout, B. H. Eussen, A. De Klein, L. Wessels, W. De Laat, et al., Domain organization of human chromosomes revealed by mapping of nuclear lamina interactions, Nature 453 (2008) 948–951.

[8] B. v. Steensel, S. Henikoff, Identification of in vivo dna targets of chromatin proteins using tethered dam methyltransferase, Nature biotechnology 18 (2000) 424–428.

[9] J. Kind, B. van Steensel, Stochastic genome-nuclear lamina interactions: modulating roles of lamin a and baf, Nucleus 5 (2014) 124–130.

[10] J. Kind, B. van Steensel, Genome–nuclear lamina interactions and gene regulation, Current opinion in cell biology 22 (2010) 320–325.

[11] B. Van Steensel, A. S. Belmont, Lamina-associated domains: links with chromosome architecture, heterochromatin, and gene repression, Cell 169 (2017) 780–791.

[12] T. R. Luperchio, M. E. Sauria, V. E. Hoskins, X. Wong, E. DeBoy, M.-C. Gaillard, P. Tsang, K. Pekrun, R. A. Ach, N. Yamada, et al., The repressive genome compartment is established early in the cell cycle before forming the lamina associated domains, BioRxiv (2018) 481598.

[13] S. Chen, T. R. Luperchio, X. Wong, E. B. Doan, A. T. Byrd, K. R. Choudhury, K. L. Reddy, M. S. Krangel, A lamina-associated domain border governs nuclear lamina interactions, transcription, and recombination of the tcrb locus, Cell reports 25 (2018) 1729–1740.

[14] J. Paulsen, M. Sekelja, A. R. Oldenburg, A. Barateau, N. Briand, E. Delbarre, A. Shah, A. L. Sørensen, C. Vigouroux, B. Buendia, P. Collas, Chrom3D: three-dimensional genome modeling from Hi-C and nuclear lamin-genome contacts, Genome Biology 18 (2017) 21.

[15] X. Lin, Y. Qi, A. P. Latham, B. Zhang, Multiscale modeling of genome organization with maximum entropy optimization, The Journal of Chemical Physics 155 (2021) 010901.

[16] A. M. Chiariello, S. Bianco, A. Esposito, L. Fiorillo, M. Conte, E. Irani, F. Musella, A. Abraham, A. Prisco, M. Nicodemi, Physical mechanisms of chromatin spatial organization, The FEBS Journal 289 (2022) 1180–1190.

[17] M. Di Pierro, B. Zhang, E. L. Aiden, P. G. Wolynes, J. N. Onuchic, Transferable model for chromosome architecture, Proceedings of the National Academy of Sciences of the United States of America 113 (2016) 12168–12173.

[18] M. Di Pierro, R. R. Cheng, E. Lieberman Aiden, P. G. Wolynes, J. N. Onuchic, De novo prediction of human chromosome structures: Epigenetic marking patterns encode genome architecture, Proceedings of the National Academy of Sciences of the United States of America 114 (2017) 12126–12131.

[19] V. G. Contessoto, R. R. Cheng, J. N. Onuchic, Uncovering the statistical physics of 3d chromosomal organization using data-driven modeling, Current Opinion in Structural Biology 75 (2022) 102418.

[20] H. Wang, J. Yang, Y. Zhang, J. Qian, J. Wang, Reconstruct high-resolution 3d genome structures for diverse cell-types using flamingo, Nature Communications 13 (2022) 2645.

[21] J. Liang, A. Perez-Rathke, Minimalistic 3D Chromatin Models: Sparse Interactions in Single Cells Drive the Chromatin Fold and Form Many-Body Units, Current Opinion in Structural Biology 71 (2021) 200–214.

[22] A. Maji, J. A. Ahmed, S. Roy, B. Chakrabarti, M. K. Mitra, A lamin-associated chromatin model for chromosome organization, Biophysical Journal 118 (2020) 3041–3050.

[23] M. Falk, Y. Feodorova, N. Naumova, M. Imakaev, B. R. Lajoie, H. Leonhardt, B. Joffe, J. Dekker, G. Fudenberg, I. Solovei, et al., Heterochromatin drives compartmentalization of inverted and conventional nuclei, Nature 570 (2019) 395–399.

[24] S. S. Lee, S. Tashiro, A. Awazu, R. Kobayashi, A new application of the phase-field method for understanding the mechanisms of nuclear architecture reorganization, Journal of mathematical biology 74 (2017) 333–354.

[25] R. Laghmach, M. Di Pierro, D. A. Potoyan, The interplay of chromatin phase separation and lamina interactions in nuclear organization, Biophysical Journal 120 (2021) 5005–5017.

[26] G. Bajpai, S. Safran, Mesoscale, long-time mixing of chromosomes and its connection to polymer dynamics, PLOS Computational Biology 19 (2023) e1011142.

[27] Q. Cheng, P. Delafrouz, J. Liang, C. Liu, J. Shen, Modeling and simulation of cell nuclear architecture reorganization process, Journal of computational physics 449 (2022) 110808.

[28] A. Yildirim, N. Hua, L. Boninsegna, Y. Zhan, G. Polles, K. Gong, S. Hao, W. Li, X. J. Zhou, F. Alber, Evaluating the role of the nuclear microenvironment in gene function by population-based modeling, Nature Structural & Molecular Biology 30 (2023) 1193–1206.

[29] L. Boninsegna, A. Yildirim, G. Polles, Y. Zhan, S. A. Quinodoz, E. H. Finn, M. Guttman, X. J. Zhou, F. Alber, Integrative genome modeling platform reveals essentiality of rare contact events in 3d genome organizations, Nature methods 19 (2022) 938–949.

[30] I. S. Tolokh, N. A. Kinney, I. V. Sharakhov, A. V. Onufriev, Strong interactions between highly-dynamic lamina-associated domains and the nuclear envelope stabilize the 3d architecture of drosophila interphase chromatin, BioRxiv (2022).

[31] F. Forsberg, A. Brunet, T. M. L. Ali, P. Collas, Interplay of lamin a and lamin b lads on the radial positioning of chromatin, Nucleus 10 (2019) 7–20.

[32] D. Peric-Hupkes, W. Meuleman, L. Pagie, S. W. Bruggeman, I. Solovei, W. Brugman, S. Gräf, P. Flicek, R. M. Kerkhoven, M. van Lohuizen, et al., Molecular maps of the reorganization of genome-nuclear lamina interactions during differentiation, Molecular cell 38 (2010) 603–613.

[33] A. Perez-Rathke, Q. Sun, B. Wang, V. Boeva, Z. Shao, J. Liang, CHROMATIX: computing the functional landscape of many-body chromatin interactions in transcriptionally active loci from deconvolved single cells, Genome Biology 21 (2020) 13. Number: 1.

[34] M. Barbieri, M. Chotalia, J. Fraser, L.-M. Lavitas, J. Dostie, A. Pombo, M. Nicodemi, Complexity of chromatin folding is captured by the strings and binders switch model, Proceedings of the National Academy of Sciences of the United States of America 109 (2012) 16173–16178.

[35] G. Gürsoy, Y. Xu, A. L. Kenter, J. Liang, Computational construction of 3D chromatin ensembles and prediction of functional interactions of alpha-globin locus from 5C data, Nucleic Acids Research 45 (2017) 11547–11558. Number: 20.

[36] Q. Sun, A. Perez-Rathke, D. M. Czajkowsky, Z. Shao, J. Liang, High-resolution single-cell 3d-models of chromatin ensembles during drosophila embryogenesis, Nature communications 12 (2021) 1–12.

[37] M. Chiang, D. Michieletto, C. A. Brackley, N. Rattanavirotkul, H. Mohammed, D. Marenduzzo, T. Chandra, Polymer Modeling Predicts Chromosome Reorganization in Senescence, Cell Reports 28 (2019) 3212–3223.e6.

[38] Y. Qi, B. Zhang, Predicting three-dimensional genome organization with chromatin states, PLoS computational biology 15 (2019) e1007024.

[39] G. Gürsoy, Y. Xu, J. Liang, Spatial organization of the budding yeast genome in the cell nucleus and identification of specific chromatin interactions from multi-chromosome constrained chromatin model, PLOS Computational Biology 13 (2017) e1005658.

[40] K. Rippe, Genome organization and function in the cell nucleus, John Wiley & Sons, 2012.

[41] D.-H. Kim, B. Li, F. Si, J. M. Phillip, D. Wirtz, S. X. Sun, Volume regulation and shape bifurcation in the cell nucleus, Journal of cell science 128 (2015) 3375–3385.

[42] G. Gürsoy, Y. Xu, A. L. Kenter, J. Liang, Spatial confinement is a major determinant of the folding landscape of human chromosomes, Nucleic Acids Research 42 (2014) 8223–8230. Number: 13.

[43] N. Briand, P. Collas, Lamina-associated domains: peripheral matters and internal affairs, Genome biology 21 (2020) 1–25.

[44] J. Kind, L. Pagie, H. Ortabozkoyun, S. Boyle, S. S. De Vries, H. Janssen, M. Amendola, L. D. Nolen, W. A. Bickmore, B. van Steensel, Single-cell dynamics of genome-nuclear lamina interactions, Cell 153 (2013) 178–192.

[45] J. Kind, L. Pagie, S. S. de Vries, L. Nahidiazar, S. S. Dey, M. Bienko, Y. Zhan, B. Lajoie, C. A. de Graaf, M. Amendola, et al., Genome-wide maps of nuclear lamina interactions in single human cells, Cell 163 (2015) 134–147.

[46] A. Perez-Rathke, Q. Sun, B. Wang, V. Boeva, Z. Shao, J. Liang, CHROMATIX: computing the functional landscape of many-body chromatin interactions in transcriptionally active loci from deconvolved single cells, Genome Biology 21 (2020) 13.

[47] S. M. Bondarenko, I. V. Sharakhov, Reorganization of the nuclear architecture in the drosophila melanogaster lamin b mutant lacking the caax box, Nucleus 11 (2020) 283–298.

[48] R. Mayer, A. Brero, J. Von Hase, T. Schroeder, T. Cremer, S. Dietzel, Common themes and cell type specific variations of higher order chromatin arrangements in the mouse, BMC cell biology 6 (2005) 1–22.

[49] Y. Benjamini, Y. Hochberg, Controlling the false discovery rate: a practical and powerful approach to multiple testing, Journal of the Royal statistical society: series B (Methodological) 57 (1995) 289–300.

[50] E. Meshorer, T. Misteli, Chromatin in pluripotent embryonic stem cells and differentiation, Nature reviews Molecular cell biology 7 (2006) 540–546.

[51] S. Heinz, C. Benner, N. Spann, E. Bertolino, Y. C. Lin, P. Laslo, J. X. Cheng, C. Murre, H. Singh, C. K. Glass, Simple combinations of lineage-determining transcription factors prime cis-regulatory elements required for macrophage and b cell identities, Molecular cell 38 (2010) 576–589.

[52] E. Panatta, A. Butera, I. Celardo, M. Leist, G. Melino, I. Amelio, p53 regulates expression of nuclear envelope components in cancer cells, Biology Direct 17 (2022) 38.

[53] M. Raices, L. Bukata, S. Sakuma, J. Borlido, L. S. Hernandez, D. O. Hart, M. A. D’Angelo, Nuclear pores regulate muscle development and maintenance by assembling a localized mef2c complex, Developmental cell 41 (2017) 540–554.

[54] X. Chen, Y. Ke, K. Wu, H. Zhao, Y. Sun, L. Gao, Z. Liu, J. Zhang, W. Tao, Z. Hou, et al., Key role for ctcf in establishing chromatin structure in human embryos, Nature 576 (2019) 306–310.

